# Reducing phenotypic instabilities of microbial population during continuous cultivation based on cell switching dynamics

**DOI:** 10.1101/2021.01.13.426484

**Authors:** Thai Minh Nguyen, Samuel Telek, Andrew Zicler, Juan Andres Martinez, Boris Zacchetti, Julian Kopp, Christoph Slouka, Christoph Herwig, Alexander Grünberger, Frank Delvigne

**Affiliations:** Terra research and teaching centre, Microbial Processes and Interactions (MiPI), Gembloux Agro-Bio Tech, University of Liège, Gembloux, Belgium; Christian Doppler Laboratory for Mechanistic and Physiological Methods for Improved Bioprocesses, Institute of Chemical, Environmental and Biological Engineering, Vienna University of Technology, 1060 Vienna, Austria; Research Division Biochemical Engineering, Institute of Chemical Environmental and Bioscience Engineering, Vienna University of Technology, Vienna, Austria; Multiscale Bioengineering, Technical Faculty, Bielefeld University, Bielefeld Germany & CeBiTec, Bielefeld University, Bielefeld, Germany

**Keywords:** Segregostat, Single cell, Flow cytometry, Phenotypic switching, Biological oscillation, Biological noise

## Abstract

Predicting the fate of a microbial population (i.e., growth, gene expression…) remains a challenge, especially when this population is exposed to very dynamic environmental conditions, such as those encountered during continuous cultivation. Indeed, the dynamic nature of continuous cultivation process implies the potential deviation of the microbial population involving genotypic and phenotypic diversification. This work has been focused on the induction of the arabinose operon in *Escherichia coli* as a model system. As a preliminary step, the GFP level triggered by an arabinose-inducible P_*araBAD*_ promoter has been tracked by flow cytometry in chemostat with glucose-arabinose co-feeding. For a large range of glucose-arabinose co-feeding, the simultaneous occurrence of GFP positive and negative subpopulation was observed. In a second set of experiments, continuous cultivation was performed by adding either glucose or arabinose, based on the ability of individual cells for switching from low GFP to high GFP states, according to a technology called segregostat. In segregostat mode of cultivation, on-line flow cytometry analysis was used for adjusting the arabinose/glucose transitions based on the phenotypic switching capabilities of the microbial population. This strategy allowed finding an appropriate arabinose pulsing frequency, leading to a prolonged maintenance of the induction level with limited impact on phenotypic diversity for more than 60 generations. This result suggests that constraining individual cells into a given phenotypic trajectory is maybe not the best strategy for directing cell population. Instead, allowing individual cells switching around a predefined threshold seems to be a robust strategy leading to oscillating, but predictable, cell population behavior.

## Introduction

Due to the inherent intracellular biological noise and under specific diversification pressures, microbial cells within the same population tend to split into subpopulations exhibiting different metabolic features [1][2][3]. This cell-to-cell variability in metabolic activities has long been recognized as an adverse effect for bioprocessing [4]. In general, it is assumed that a highly homogeneous microbial population leads to more tractable, robust and predictable population behaviour [5][6]. However, in nature microbial cells have evolved to constantly adapt to different diversification pressures and phenotypic plasticity is known to improve cellular decision-making processes, resulting in most of the cases in a fitness gain for the whole population [7]. Adaptation to carbon limitation [8] and switching to an alternative carbon source [9][10] have been found to be key drivers of microbial phenotypic diversification. Optimal exploitation of the functionalities offered by this diversification process by microbial populations typically led to improved fitness in front of environmental perturbations [11], notably through mechanisms such as bet-hedging [12][13]. Real occurrence of bet-hedging in cell population is still a debate, but such phenomenon has been observed to occur from relevant gene circuits driving e.g., the switch to alternative carbon sources in bacteria [10][12] or the starvation response in yeast [14][15]. Bet-hedging relies on the simultaneous occurrence of subpopulations of cells pre-adapted to diverse environmental conditions, leading to an anticipative adaptation to unexpected environmental transition [16]. It has been mathematically demonstrated that this strategy is particularly beneficial in fluctuating environments with given fluctuation frequencies and amplitudes [17]. More specifically, high phenotypic diversity has been found to lead to a gain in fitness if the rate of phenotypic switching is equal or lower than the rate of environmental transitions. Phenotypic switching dynamics are complex and involve several layers of regulation, each at specific time-scales, i.e. transcription or translation, noise propagation through the gene regulatory network (GRN) [18] and possible feedback effects exerted by the cell division process, resulting in the dilution over the intracellular molecular species [19]. These switching mechanisms are then dependent on the specifics of the biological system and its culturing processing conditions, therefore they must be explored and analyzed based on single cell data. In this work, population stability in a continuous cultivation was controlled via population heterogeneity measurements coupled with tailored carbon source feeding strategies. In this context, the induction of the arabinose operon in *Escherichia coli* following glucose scarcity and switch to arabinose utilization was used as a tunable model system. The strategy relying on the use of environmental perturbations at given frequencies and amplitudes for stabilizing and directing microbial diversification processes was explored through an experimental device previously set-up and published, i.e. the segregostat [20]. The segregostat is a continuous cultivation device connected to on-line flow cytometry for tracking phenotypic diversification processes. More importantly, the device is able to automatically induce environmental transitions based on the phenotypic switching ability of the population. However, the idea developed in this work is to allow cells freely switching around a predefined fluorescence threshold, instead of constraining cells into a predefined fluorescence time course as shown in previous work aiming at controlling gene expression in cell population [21][22][23]. In this work, this device was used to assess whether stochastic switching could be applied for stabilizing microbial populations under continuous cultivation. Surprisingly, when setting this switching rate for maximizing phenotypic diversification dynamics, the segregostat led to a more predictable induction profile of the arabinose-inducible promoter (P_*araBAD*_), which remained stable over more than 60 cell generations. In comparison to chemostat cultivation, where subpopulations of cells with different GFP content was observed, the segregostat led to a uniform population of cells smoothly transitioning between the uninduced (low GFP) and induced (high GFP) states.

## Material and Methods

### Strains and media

*E. coli K12 W3110* wildtype was transformed with pBbB8k (4480 bp) plasmid with the P_*araBAD*_::GFP, which is an arabinose inducible promoter system. This plasmid belongs to a library of expression vectors compatible with the BglBrick standard [24]. Pre-cultures and cultures were performed on defined mineral salt medium containing (in g.L^−1^): K_2_HPO_4_ 14.6, NaH_2_PO_4_.2H_2_O 3.6; Na_2_SO_4_ 2; (NH_4_)_2_SO_4_ 2.47, NH_4_Cl 0.5, (NH_4_)_2_-H-citrate 1, glucose 5, thiamine 0.01. Thiamine was sterilized by filtration (0.2 μm). The medium was supplemented with 3 ml l^−1^ of trace element solution, 3 ml l^−1^ of a FeCl_3_.6H_2_O solution (16.7 g.L^−1^), 3 ml l^−1^ of an EDTA solution (20.1 g l^−1^) and 2 ml.L^−1^ of a MgSO_4_ solution (120 g.L^−1^). The trace element solution contained (in g.L^−1^): CaCl_2_.H_2_O 0.74, ZnSO_4_.7H_2_O 0.18, MnSO_4_.H_2_O 0.1, CuSO4.5H_2_O 0.1, CoSO_4_.7H_2_O 0.21. Kanamycin (50 μg/mL) was used as a selective marker.

### Chemostat cultivations

Chemostat cultures were performed in a 2MAG© block systems (2mag AG, Munich, Germany). The bioreactor system was equipped with positions for eight fermentation vessels with 16 mL total volume and working volume of 10 mL. The bioreactor block was equipped with magnetic inductive drives with two independent heat exchangers integrated into the bioreactor block, one for temperature control for the reaction broth and the second to control the headspace temperature and prevent evaporation. The system was also equipped with fluorometric sensor spots for dissolved oxygen (DO) positioned at the bottom of each reactor (minireaktor HTBD, Presens, Regensburg, Germany). Each reactor was equipped with a magnetic S-impeller with two permanent magnets (Sm2Co17, IBS magnet, Berlin Germany). In each chemostat, fresh medium was pumped into the culturing chamber at a constant rate, while culture effluent exits at an equal rate though the inlet and outlet tubes with an inner diameter of 0.8 mm (Marprene tubing; Watson-Marlow, Rommerskirchen, Germany). Overnight pre-cultures w performed in 1L baffled flasks containing 100 ml of culture medium and stirred with 200 rpm at 37°C. Fermenter inoculation was adjusted to 0.5 OD_600_ for the initial batch phase. The temperature was maintained at 37°C under continuous stirring rate of 2600 rpm. The DO signal was used as an indicator for switching to the continuous operation modes (a rapid increase of the DO signals after depletion of glucose in the initial batch medium, observed typically after 3–5 h). The medium was continuously fed with the complete minimal medium containing glucose and arabinose at different ratios at a dilution rate of 0.5 h^−1^. Off-line samples were taken from the reactors each 5-10 h and the analysis of the GFP expression levels was performed with a BD Accuri C6 (BD Biosciences, CA, USA) based on FL1-A channel (excitation 488 nm, emission 533 nm).

### Segregostat cultivations

Cultures in segregostat mode were performed in lab-scale stirred bioreactor (Biostat B-Twin, Sartorius) with 1 L working volume. The batch phase was started at an OD_600_ value of 0.5. The pH was maintained at 7.0 by automatic addition of ammonia solution 25%(w/v) or phosphoric acid 25% (w/v). The temperature was maintained at 37°C under a continuous stirring rate of 800 rpm and an aeration rate of 1 vvm. The DO signal was used as an indicator for switching to the continuous operation modes (a rapid increase of the DO signals after depletion of glucose in the initial batch medium, observed typically after 3–5 h). The feeding medium, containing 5 g/L glucose, was continuously fed with a dilution rate of 0.5 h^−1^. According to the sequences controlled by the online FC software, pulse of arabinose was injected in order to quickly increase (approx. 30 seconds) the global arabinose concentration to 0.5 g/L [20]. On-line sampling was performed every 12 minutes and processed according to the dilution sequence set through the following series of steps. First, the sample is automatically transferred to C6 FC (BD Accuri C6, BD Biosciences) and analyzed at a medium flow rate (33 μl min^−1^). All the data related to the different parameters (mean, median, CV) are displayed in real-time during the cultivation. Then a tailor-made feedback control loop MATLAB script based on the FC data regulated the actuator profile for the arabinose addition. Within this script, FC events were gated based on forward scatter (FSC) to distinguish to single cells and clumps. Then GFP positive events were gated based on a fluorescence threshold value of 1000 (arbitrary units, FL1-A; excitation 488 nm, emission 533 nm) for the first segregostat (considered as a loose control policy) and with a threshold fluorescence value of 10000 for the second segregostat (considered as a tight control policy). Control policy was set for actuator activation when the fraction of GFP positive cells was measured to be below 50%. Actuator activation then exerted arabinose pulsing by the use of a digital control system comprising a peristaltic pump (Watson Marlow, 101 UR). Each segregostat experiment were carried out in duplicate. Data were exported as .fcs files and processed by a custom Python script (see next sections for a description of the data processing steps).

### Online flow cytometry (FC) analyses

The online flow cytometer platform comprises three modules and can be operated in segregostat mode: (i) a conventional culture device, (ii) a physical interface for sampling and dilution containing peristaltic pumps and mixing chamber, and (iii) a detection device, in this work an Accuri C6 flow cytometer (BD Accuri, San Jose CA, USA) as used. Briefly, sample processing comprises the following steps: (i) sample acquisition, (ii) online FC analysis, (iii) dilution threshold and (iv) feedback control loop. The sample is entered and removed from the mixing chamber based on silicone tubing (internal diameter: 0.5 mm; external diameter: 1.6 mm, VWR, Belgium) and peristaltic pumps (400FD/A1 OEM-pump ~13 rpm and 290 rpm, Watson Marlow). Before and after each experiment, all the connection parts (tubing, pumps and mixed chamber) were cleaned with ethanol and rinsed with filtered PBS as set-up in a previous protocol [25].

### On-line FC data processing and accessibility

Segregostat experiment involves the generation of a huge amount of FC data. A typical run of 100 hours led to 500 FC analyses, with a total of approximately 10^7^ cells processed. Before processing, these independent .fcs files were compacted in a dataframe (.pkl file extension) based on the Pandas package (https://pandas.pydata.org/) from Python. The codes for generating the .pkl files and the figures are available on a GitLab repository https://gitlab.uliege.be/F.Delvigne/paper_segregostat_arabinose. The code with name “From_fcs_to_pkl_and_statistics.py” can be used for generating most of the figures presented in this paper (i.e., figure 3 and 5). The code with name “Plots_from_pkl.py” can be used for generating the FC dotplots from a .pkl file and for each time points. These dotplots were further assembled into a single .avi movie file based on the ImageJ software [26] shown as Supplemental material (Movies S1 and S2). Raw .fcs data were deposited on FlowRepository and can be accessed by following the links below:

- Segregostat with low fluorescence threshold (1000 FU, loose control policy), replicate 1:
- Segregostat with low fluorescence threshold (1000 FU, loose control policy), replicate 2:
- Segregostat with high fluorescence threshold (10000 FU, tight control policy), replicate 1:
- Segregostat with high fluorescence threshold (10000 FU, tight control policy), replicate 2:

### Metabolite analysis

Supernatant glucose and arabinose concentrations were analyzed by high-performance liquid chromatography (Waters Acquity UPLC^®^ H-Class System) using an ion-exchange Aminex HPX-87H column (7.8 × 300 mm, Bio-Rad Laboratories N.V.). The analysis was carried out with an isocratic flow rate of 0.6 mL min^−1^ for 25 min at 50°C. The mobile phase was composed of an aqueous solution of 5 mM H_2_SO_4_. Elution profiles were monitored through a Waters Acquity^®^ Refractive Index Detector (RID) (Waters, Zellik, Belgium). Glucose and arabinose standard solutions (Sigma-Aldrich, Overijse, Belgium) were used to determine the retention times and construct calibration curves.

### Transcripts isolation and quantification

Samples were taken at different times during fermentation. Samples were immediately frozen and kept at − 80°C. The total RNA was extracted using the NucleoSpin RNA Mini kit (Macherey-Nagel, Germany) following the manufacturer’s protocols. qPCRs measurements were conducted in an Applied Biosystems Step One Real-Time PCR system (Life Technologies, Grand Island, NY, USA). The thermal cycling protocol was performed as follows: initial denaturation for 15 min at 95°C, 40 cycles of 95°C for 10 sec, at 65°C for 15 sec, and at 72°C for 20 sec, and a final dissociation cycle at 95°C for 30 sec, 65°C. qPCR data was normalized to the housing keeping gene *ihfB,* and the sample taken from the end of batch phase was used as a reference [27]. The 2(-Delta Delta C(T)) method was used to process the data and compute the relative changes in gene expression [28].

## Results

### Chemostat with glucose/arabinose co-feeding leads to the formation of subpopulations of cells exhibiting different GFP content

In a first set of experiments, classical chemostat cultivation has been run at a dilution rate of 0.5 h^−1^. Combinations of glucose and arabinose were considered as co-feeding strategy for activating the arabinose operon during the continuous cultivation phase, and the population heterogeneity was determined based on the signal provided by the GFP reporter system expressed by the means of the P_*araBAD*_ promoter (Figure 1A). Flow cytometry analysis revealed that the co-feeding strategy with arabinose and glucose led to a heterogeneous GFP expression profile, traduced by the appearance of two subpopulation of cells (Figure 1B). Indeed, the simultaneous presence of GFP negative and GFP positive subpopulations was observed right after the induction upon activation of the co-feeding regime. This “all-or-none” response has previously been observed under similar operating conditions with arabinose/glucose [29] and lactose/glucose co-feeding [30]. This behavior was observed for different arabinose/glucose ratios (Figure 1C and 1D).

**Figure 1:**
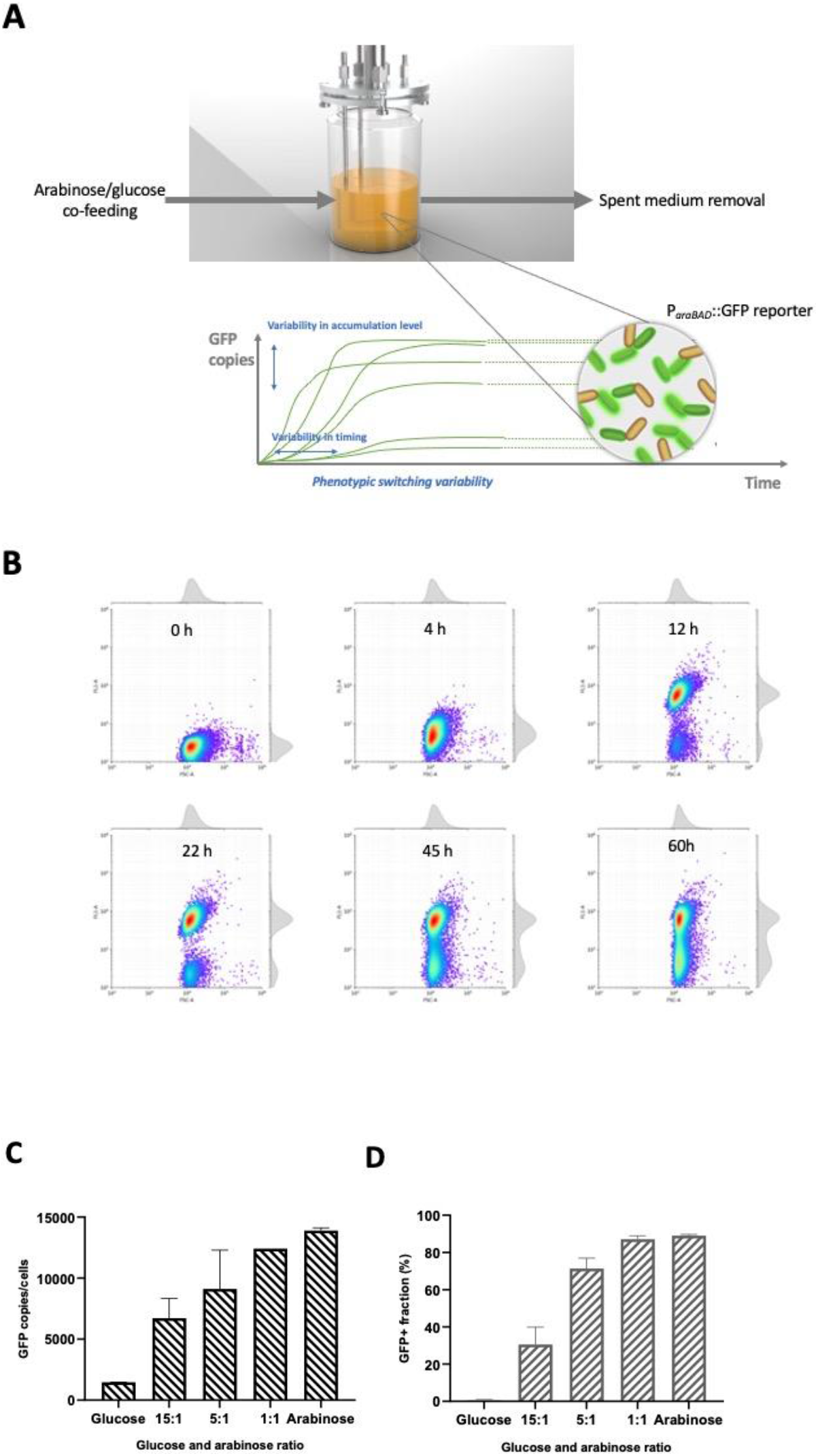
**A** Sketch of the chemostat set-up used for the glucose-arabinose co-feeding experiments. **B** Flow cytometry monitoring (x-axis: forward scatter signal; y-axis: green fluorescence signal accounting for the accumulation of GFP inside cells) of a chemostat with glucose-arabinose co-feeding (dilution rate 0.5 h-1; ration glucose-arabinose 5:1). **C** Evolution of the GFP positive cell fraction and **D** the mean GFP copies/cell

### Design of a control strategy based on cell phenotypic switching dynamics

Based on the data gathered from chemostat with glucose-arabinose co-feeding, it is clear that a classical control procedure based on the definition of a setpoint (i.e., the quantity of GFP accumulated in cells in our case) and the use of a Proportional-Integral-Differential (PID) controller is not suitable (Figure 2A). Indeed, considering the diversity of the GFP values observed during continuous culture, the proper identification of a given setpoint is impossible. The picture is even more complicated if the simultaneous occurrence of GFP negative and GFP positive subpopulations is considered. Previous researches have pointed out that PID control is not appropriate for controlling synthetic gene circuits within cell population [31]. As an example, Lugagne and co-workers have demonstrated that bacterial population engineered with a synthetic gene circuit can be stabilized based on simple ON/OFF control strategy involving the periodic addition of inducers [32]. Since biological noise is impairing the efficient utilization of a classical PID strategy for directing cell population, a new approach is then needed. Whereas sophisticated control procedure relying on model-predictive approaches have already been investigated [33][23], we decided to consider a simple control procedure based on the intrinsic switching properties of microbial cells. Indeed, instead of constraining cell fluorescence trajectories around a given setpoint, we allowed them to freely switch around a predefined value. The critical step remains the appropriate identification of the fluorescence threshold fluorescence value to be used for defining the cell switching process. As stated before, two subpopulations of cells, differing in global GFP content, can be observed i.e., GFP negative (“low” state) and GFP positive (“high” state), and we focused on this specific population structure for defining the switching thresholds (Figure 2A). A cell-machine interface relying on the on-line FC monitoring of the P_*araBAD*_::*GFP* reporter was designed for automatically triggering the addition of arabinose pulses during continuous cultivation based on the phenotypic switching capabilities of microbial cells [20]. It has been shown previously that microbial cells are able to switch stochastically from one phenotypic state to another in response to environmental perturbations [17]. In the present work, this specific behavioral feature was exploited for optimizing the GFP expression profiles for the whole population. The cell-machine interface builds up on the previously described segregostat concept. This system, relying on the use of feedback control based on on-line FC measurement, has been previously used for minimizing phenotypic heterogeneity related to the membrane permeabilization process during *E. coli* continuous cultivation [20][34]. In its current version, the segregostat comprises a continuous cultivation device connected to an on-line flow cytometry device allowing sampling the state of the population at regular time intervals. This device then allows optimizing the glucose-arabinose switches based on the actual phenotypic switching dynamics of the microbial population. According to the bimodal behavior of the cell population observed during chemostat cultivation (Figure 2A), two different thresholds have been considered (Figure 2B and 2C).

**Figure 2:**
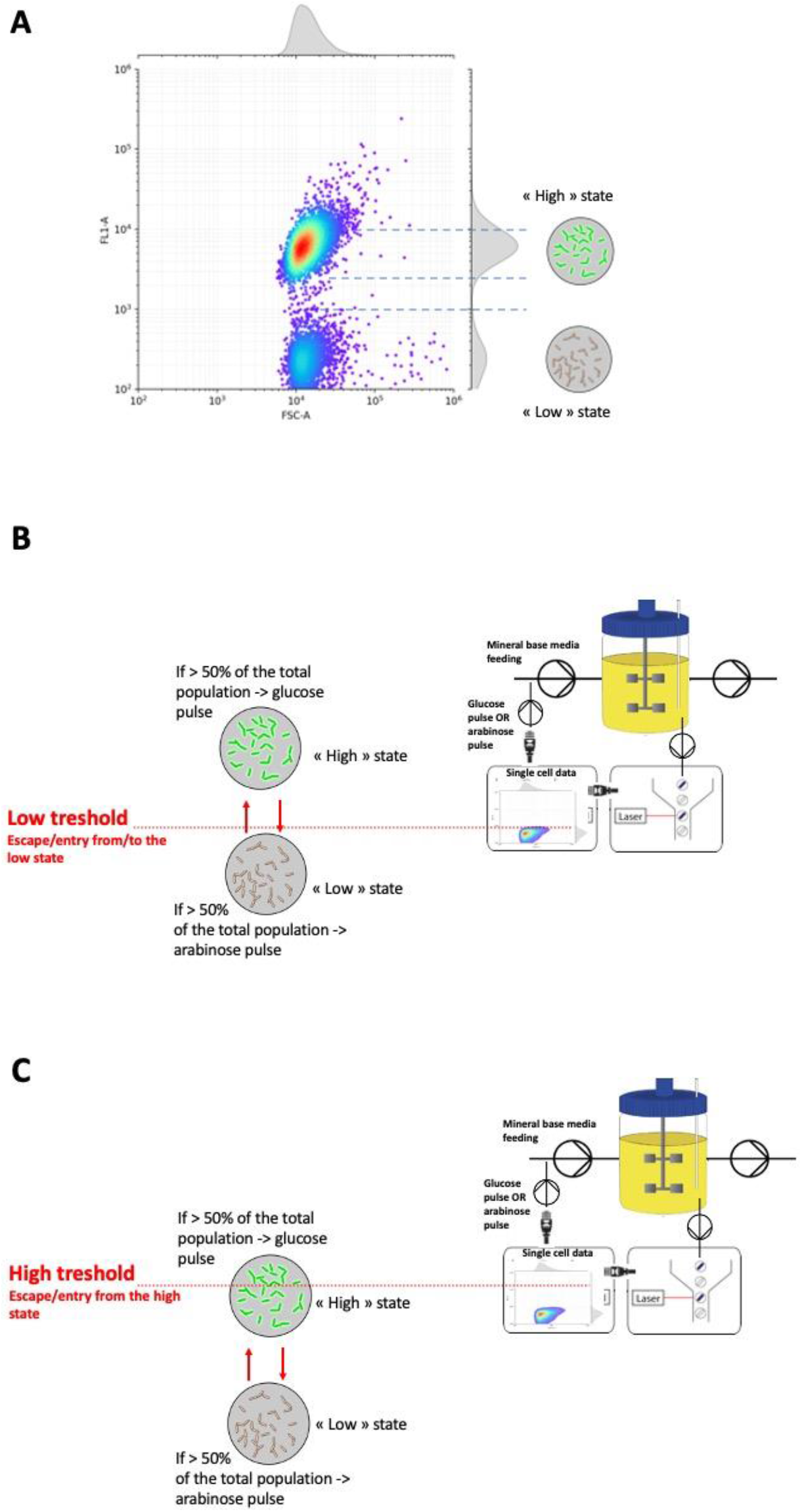
**A** Design of a population control strategy based on cell phenotypic switching dynamics and on the bimodal GFP distribution. **B** Adjustment of arabinose pulsing based on the phenotypic switching ability of cells. In this case, a low threshold has been selected (loose control procedure). **C** Adjustment of glucose and arabinose pulsing based on the phenotypic switching ability of cells. In this case a high threshold has been selected (tight control procedure).

### Automated addition of arabinose pulses based on the phenotypic switching capability of cell population leads to oscillating but stable gene expression profiles

In the first segregostat experiments, low fluorescence threshold (1000 RFU), corresponding to the upper limit of the low state (Figure 2B) was chosen. When at least 50% of the population has crossed this threshold, then an environmental fluctuation, here represented by an arabinose pulse, was triggered. According to this control policy, it was possible to maintain the whole population fluctuating between the high and the low state (Figure 3). Additionally, no bimodal GFP distribution was observed, the whole population following the glucose to arabinose transitions by dynamically adapting gene expression (Supplemental Material, Movie S1). Additionally, gene expression levels were monitored the cultivation by RT-qPCR, and residual arabinose and glucose was determined by HPLC (Supplemental Material, Figure S1). In a second step, the transition rate of cells between the low and the high state have been analyzed. It was assumed that cell stochastic switching could be modelled by a simple Poisson process. The transitions from the low to the high state (Figure 4A) and from the high to the low state (Figure 4B) were then modelled and compared to the experimental data. A transition rate of 2.8 h^−1^ was determined for the switch to the low to the high state, and the process was nicely fitted by a simple Poisson process (Figure 4A). On the other hand, a much slower transition rate was determined for the switch from the high to the low state. In this case, two phases can be observed at the level of the GFP decay plot, each of them being approximated by two Poisson processes occurring at different rate i.e., 0.02 h^−1^ and 0.3 h^−1^ respectively. These results suggests that several subpopulations of cells were involved in the decay of GFP fluorescence. Since the GFP variant used in this work was quite stable, dilution of GFP occurred mainly by cell division. Then, a cell-to-cell difference in growth rate could explain this wider heterogeneity at the level of the switching dynamics during the relaxation to the low state. Such differences have been previously reported for cells involved in diauxic shift [12][9]. It is then interesting to wonder if the segregostat could be used for further improving the GFP expression level in presence of this extra layer of biological noise. A more constraining control procedure has been considered accordingly and will be described in the next section.

**Figure 3:**
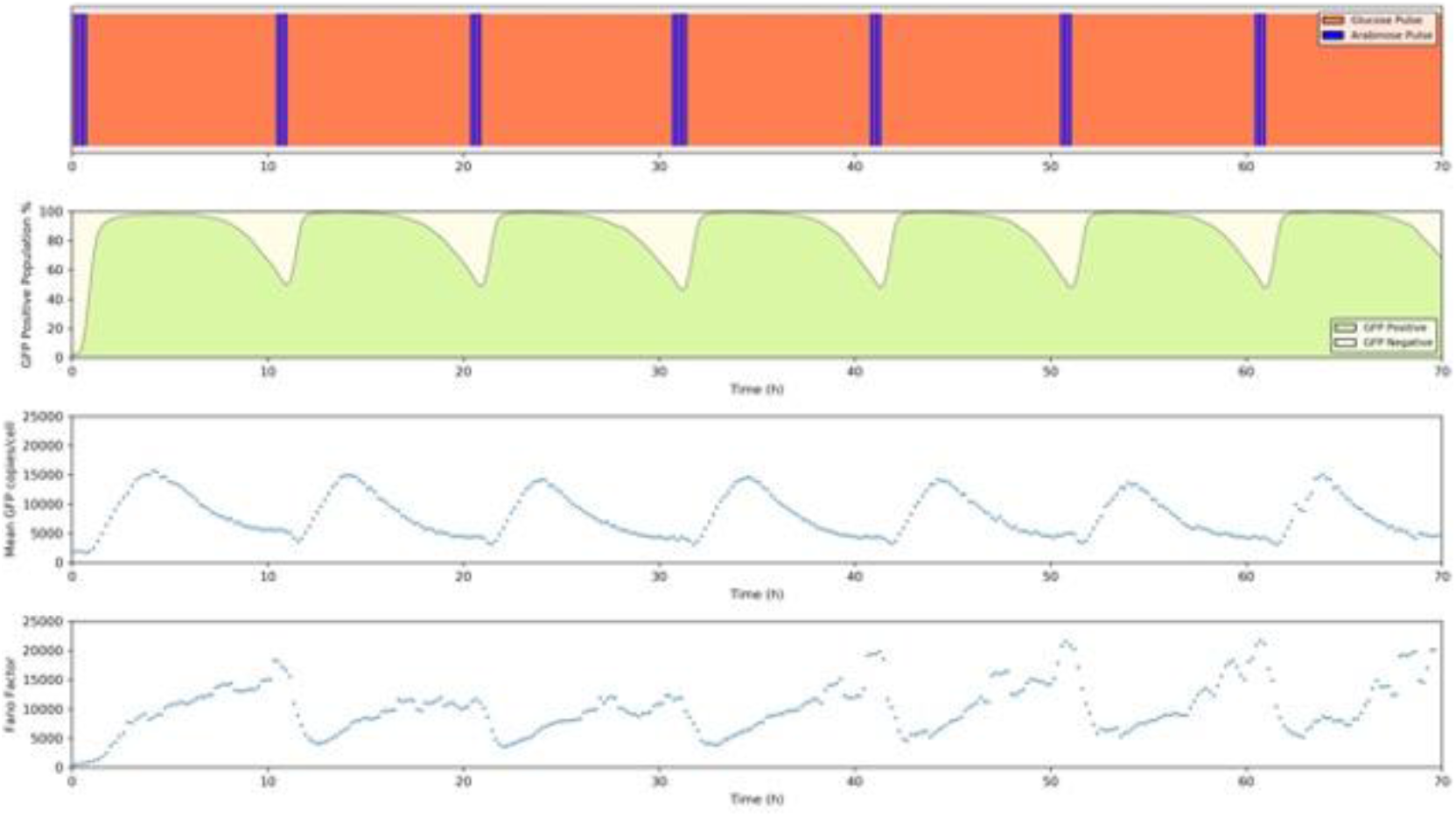
Population dynamics in function of the automated addition of glucose/arabinose pulses. Mean GFP copies per cells and Fano factor (ratio between the variance and the mean of GFP distribution) have been computed based on the GFP positive fraction only.

**Figure 4:**
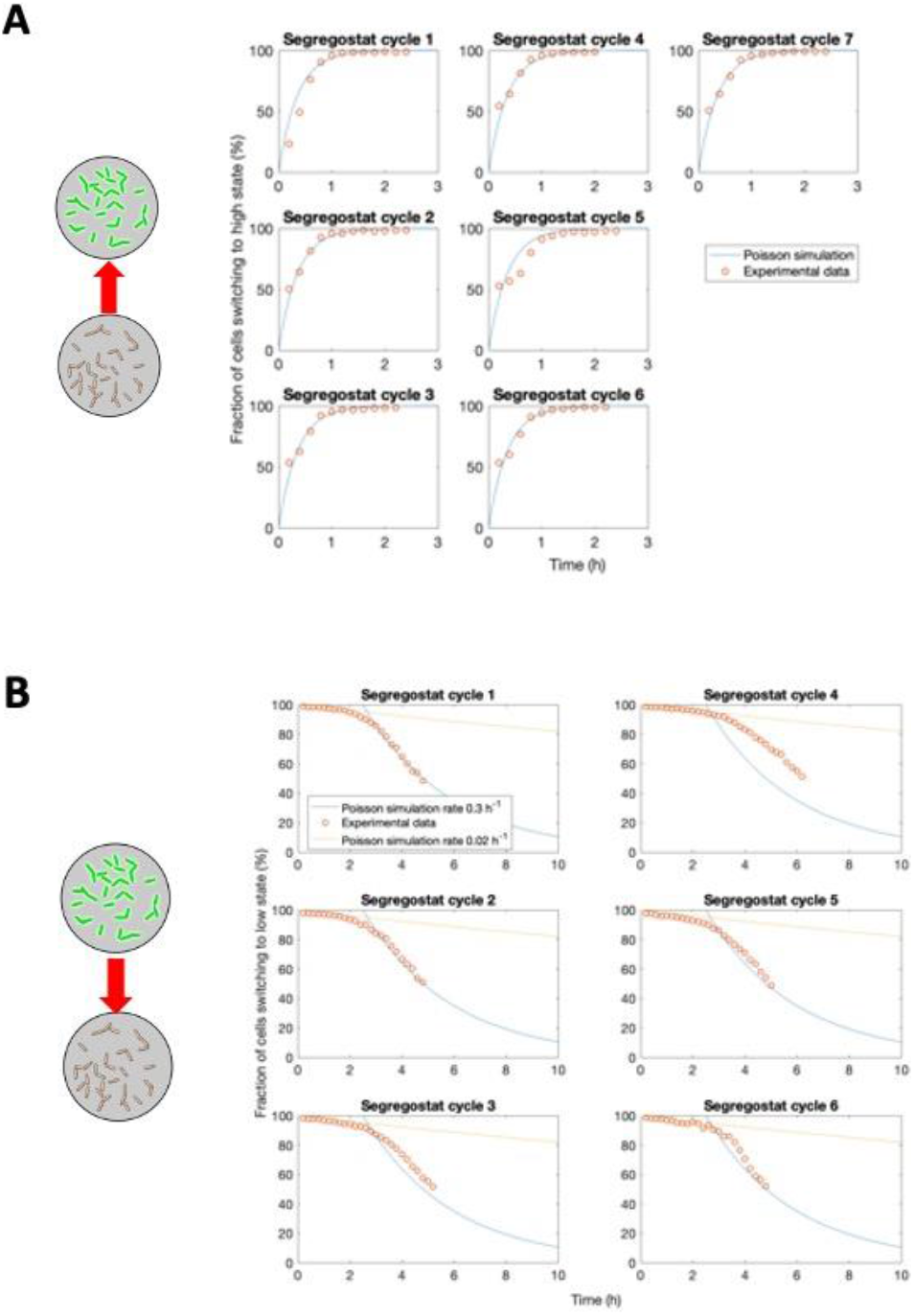
Analysis of the cell switching dynamics during segregostat experiment described in Figure 3. **A** Comparison of the experimental switching data for the transition from the low to the high state with a Poisson process. **B** Comparison of the experimental switching data for the relaxation from the high to the low state with two Poisson processes.

### Forcing the entry into the high state leads to harmonic oscillations at the level of the GFP content

For optimizing gene expression, the relaxation of cells into the low state must be prevented. Then, it would be interesting to know if it is possible to maintain the whole cell population into the high state under continuous cultivation. In order to optimize GFP accumulation, a high threshold was considered for running a second segregostat based on a more constraining control policy (Figure 2C). This threshold was set based on the upper limit of the high state (10000 RFU) and is then exerting more pressure on the cells by comparison with the experiment carried out with a loose control policy (i.e., the first segregostat with a fluorescence threshold set at 1000 RFU). Based on this new threshold, all cells were kept into the high state during the whole cultivation, with a GFP copies/cell oscillating around a higher value by comparison to what was obtained based on the loose control policy (Figure 5). Again, it is important to point out that no bimodal GFP distribution was observed during continuous cultivation under segregostat regime (Supplemental Material, Movie S2). Furthermore, these results pointed out that it is also possible to the entire cell population into the high state over long periods of time (more than 60 generations). Enhancing the switching threshold led to a transition from relaxed GFP oscillations to harmonic oscillations. Such effect was also observed when optimizing the dynamics of a synthetic oscillatory circuits [35]. Additionally, the amplitude of the oscillations was decreased and the frequency was increased, further improving the absolute gene expression of the population. Another important feature to be considered is the stability of the population. In order to analyze the robustness of the GFP oscillations obtained, correlation between the GFP level of expression (Figure 6B) and the resulting changes in cell density (Figure 6A). The resulting phase plane analysis showed that robust cycles were achieved after a couple of hours of cultivation and that a limit cycle was reached (Figure 6C). Further experiments were also performed in order to explore potential sources for these biological oscillations. Basically, biological oscillations can be generated from a system exhibiting two features i.e., a negative feedback and a delay [36][37]. In our case, negative feedback is provided by two regulatory elements in the GRN responsible for the regulation of the arabinose operon i.e., the transcriptional factors CRP and AraC. For the optimal induction of the arabinose operon the respective factors must bind to cAMP (signal for glucose limiting conditions) and arabinose (signal for the presence of arabinose). In segregostat, glucose is always maintained at a very low concentration and arabinose is pulsed according to the state of the population (Figure 7A), generating a sequential switching of cells between the low and the high state. Negative feedback leading to oscillating GFP expression pattern arises upon arabinose consumption. This consumption arises with a delay (i.e., time needed by cell population for consuming the arabinose), further strengthening the oscillating pattern. The expression level of a set of key genes were monitored during segregostat cultivation, with samples taken at the same time than the one used for determining the extracellular glucose and arabinose concentrations (Figure 7B). We can see that among the arabinose operon, the *araC* gene expression remains at the same level, whereas the *araB* genes, coding for one of the enzymes implied in arabinose utilization, fluctuates according to the arabinose pulsing. These results suggest that the genetic oscillations are due to the arabinose availability. However, the RT-qPCR data pointed out thar *crp* gene fluctuates also a lot, adding another layer of complexity to the induction mechanisms linked to the arabinose operon. The other genes tracked i.e., *acs* (implied in acetate re-utilization), *ptsG* (coding for the main glucose transporter) and *pta* (also implied in acetate re-utilization, showed no significant fluctuations following glucose-arabinose transitions.

**Figure 5:**
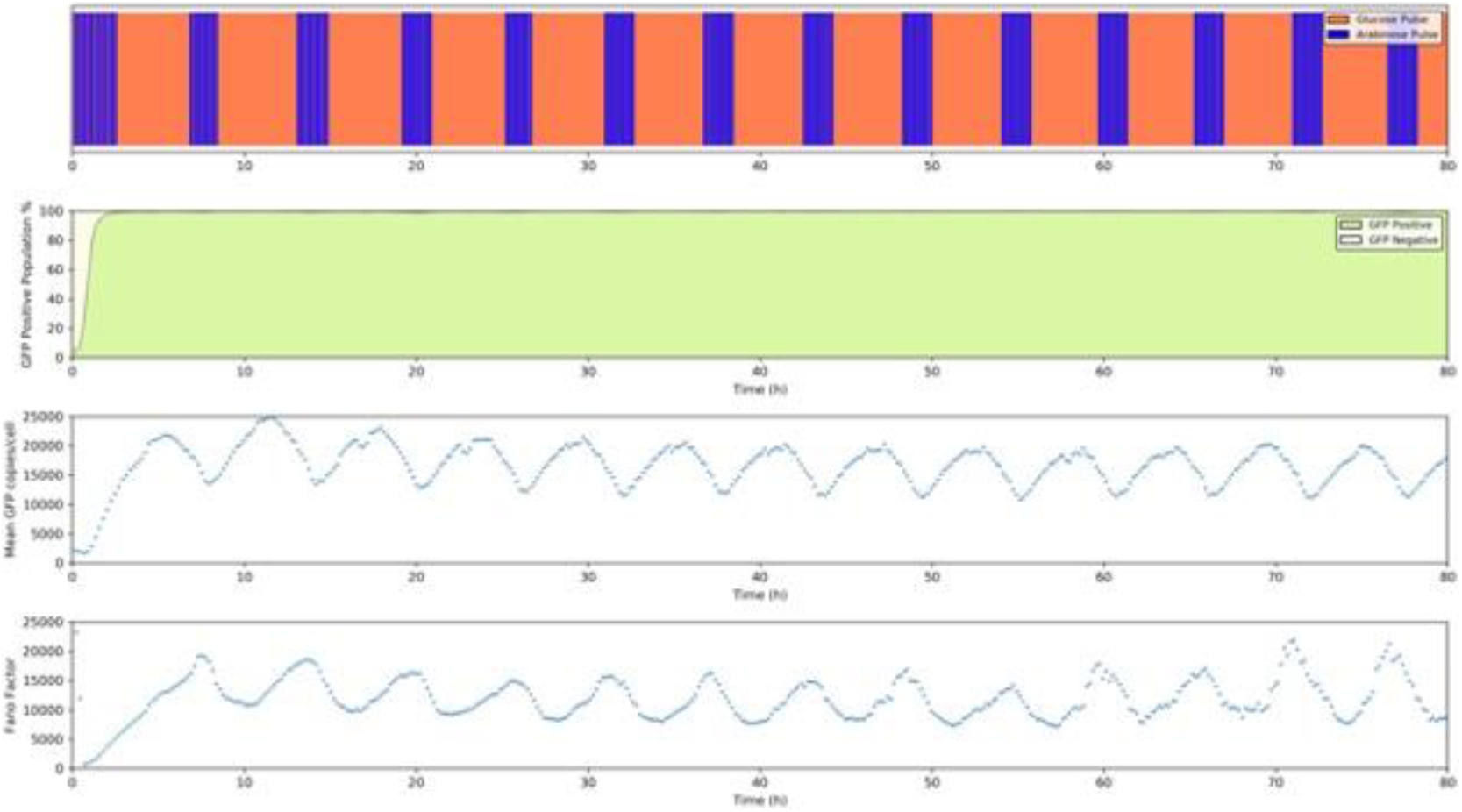
Population dynamics in function of the automated addition of arabinose pulses. Mean GFP copies per cells and Fano factor (ratio between the variance and the mean of GFP distribution) have been computed based on the GFP positive fraction only.

**Figure 6:**
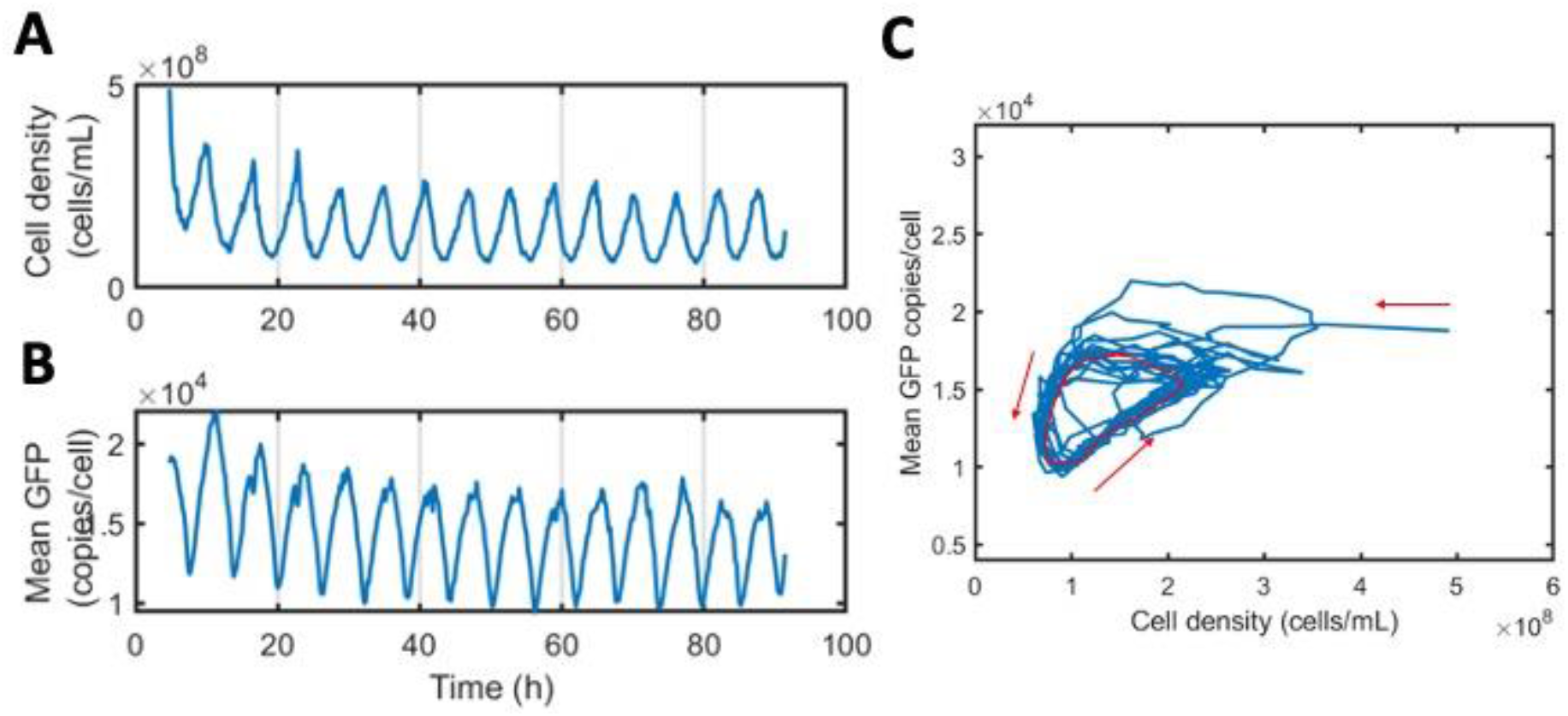
Time course of **A** the mean GFP copies/cell and **B** the cell density during segregostat experiment depicted in Figure 5**. C** phase plane analysis of the oscillations between cell density and GFP content during segregostat experiment. Limit cycle is highlighted in red.

**Figure 7:**
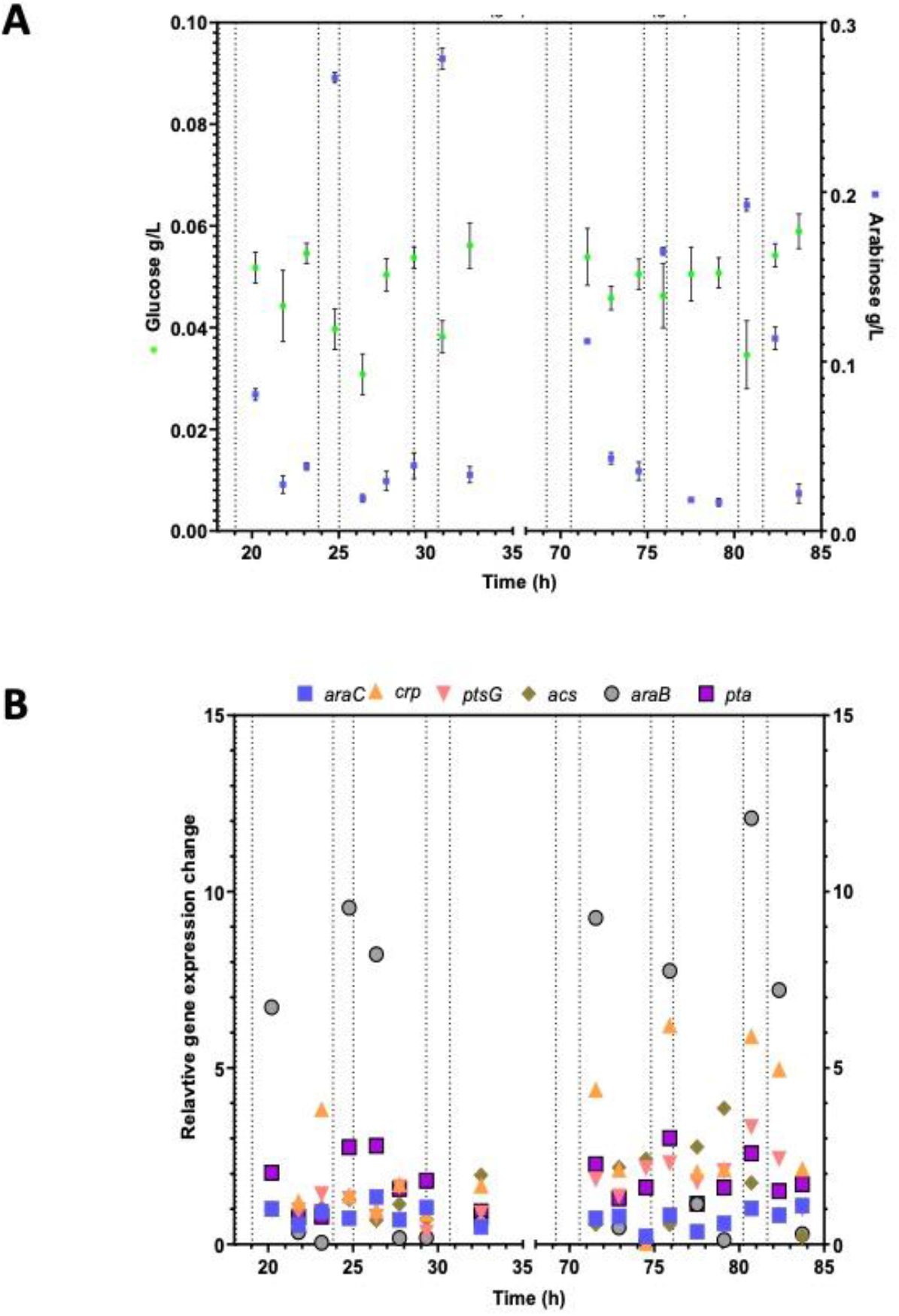
**A** Evolution of residual glucose (⚫) and arabinose (◼) concentrations for the segregostat cultivation performed based on tight control procedure (experimental settings displayed in Figure 2C). Dashed lines correspond to the arabinose pulsing phases. Results were expressed as the mean of three technical replicates. B Dynamics of the expression level of *araC* (◼), *crp*(▲), *ptsG*(▼), *acs*(◆), *araB*(⚫) and *pta*(◼) for the segregostat cultivation performed based on tight control procedure. Dashed lines correspond to the arabinose pulsing phases. Results were expressed as the mean of three technical replicates.

## Discussion

Only a few studies are actually dealing with the response of cell population to repetitive stimuli and new insights are needed at this level. The main issue is that the whole spectrum of amplitude-frequency related to environmental perturbation is too broad to be screened in a time efficient way. One of the remarkable features of the segregostat is that the environmental perturbation is triggered by the capacity of the microbial population to split into different phenotypic states, leading to the systematic determination of the optimal rate of environmental perturbations to be tested among the whole spectrum of amplitude/frequency. The use of the segregostat platform shed new light about the impact of phenotypic heterogeneity on the dynamics and stability of microbial populations. In comparison to bimodality in GFP distribution observed in chemostat with glucose/arabinose co-feeding, cultivation performed in the segregostat mode led to an unimodal GFP distribution with the microbial population smoothly transiting between the high (i.e., high GFP content driven by P_*araBAD*_ promoter) and low induction states (See movies S1 and S2 supplied in Supplemental Material for an animated evolution of GFP distribution with time). This led to a fully predictable oscillating pattern for GFP expression that was stable during the whole continuous cultivation comprising several generations. Two specific features need to be further discussed, the first one being about the way that population control is achieved and the second is about the biological oscillations observed under such control procedure. About the first feature, it is important to recall that segregostat operates by letting individual cells among the population to freely switch around a predetermined fluorescence threshold. Microbial cell among a given population naturally switch for adapting to changing environmental conditions and stresses. This is particularly true for microbial population growing under continuous cultivation conditions, where stress pressure is high and is known to force cells into different phenotypic trajectories. Accordingly, it is utopic to think that cell trajectories in gene expression can be precisely constrained [22]. Instead, it has been proposed in this work to give the opportunity to cell for switching around given gene expression threshold. This remarkable feature is quite original by comparison to the well adopted predictive model control loops generally used for directing cell populations [21][23][33]. This control strategy led to genetic oscillations that was found to be particularly robust. The second feature is about these resulting biological oscillations. It is quite surprising to obtain such oscillations, since biological oscillators are recognized for their specific architecture i.e., either gene circuits containing three mutually repressing genes (known as the repressilator)[35][38], or two genes involving the presence of a negative and a positive feedback loop [39]. However, a previous study reported that simplified circuit design could also lead to robust oscillations if stimulated at the right frequency of inducer inputs [37]. In our case, the arabinose operon is under the control of a gene circuit exhibiting a specific feedforward loop architecture [36][40]. Being cultivated under glucose limiting conditions, cell population is readily activated upon addition of the inducer arabinose. This feature has been confirmed by approximating the segregostat cycle involving the switch from the high to the low state by a simple Poisson process (Figure 4A). Biological oscillations are then generated by the negative regulation arising from the progressive disappearance of the arabinose from the cultivation medium. This disappearance occurs with a delay in time i.e., in our case the time needed by the cells to consume the arabinose, further strengthening the oscillatory behavior of the system [36][37]. In our case, phase resetting (switching from the high the low state) has indeed been shown to be dependent on cell division (Figure 4B). It has been previously demonstrated analytically that for a population of biological oscillator under entrainment, phase resetting can be achieved through cell division [41]. In our case, it is indeed suspected that cell division is implied into the relaxation from the high to the low state, with at least two subpopulation of cells exhibiting different growth rate.

Besides its utilization as an effective population control device, segregostat data can also be used for understanding the dynamics of key biological processes involved in phenotypic switching dynamics and microbial population stability. Biological noise is known to play a significant role in the process of adaptation to environmental perturbations [17][18][42], such as the switch from glucose to arabinose considered in this study. As stated in the introduction section, biological noise is quite well characterized for a single gene [43], but the picture becomes more complex in the case of complex GRN. In this case, it is particularly difficult to predict how noise propagate from a gene to another in the regulatory network [44]. The arabinose operon is under the control of a feedforward motif, allowing the cell to tightly regulate the expression of the genes needed for efficient arabinose utilization [45]. This tight regulation mechanism has been selected in order to mitigate the metabolic burden associated with the expression of the *araBADEFGH* genes by optimally inducing this operon when glucose is absent and arabinose becomes available in the environment [36]. It is also known that, under specific environmental conditions, this motif can lead to bimodal behavior with cells inducing the operon whereas others in the same population remains at a low state. This strategy could be interpreted as a bet-hedging mechanisms where microbial population effectively exploit noise in order to optimize its fitness in uncertain environmental conditions [16]. Bimodality in GFP expression has been observed in our case during chemostat experiment with glucose-arabinose co-feeding. Interestingly, this bimodal behavior can be avoided when pulsing glucose and arabinose at a particular frequency, which is in accordance with the phenotypic switching ability of cells. In this case indeed, only a unimodal GFP expressing population was observed. However, this population also exhibited noise but with a mean GFP expression level oscillating at a frequency corresponding to the environmental fluctuations. This specific mode of diversification, called dynamic heterogeneity, has been previously predicted based on mathematical modelling by Thattai and van Oudenaarden [17]. The results presented in this work point out that cell physiology in segregostat is significantly different from the one that is observed in chemostat, involving a fundamentally different modes of diversification dynamics. Further work is needed for characterizing this physiology and the underlying mechanisms behind dynamic heterogeneity.

## Conclusion

The segregostat can be used for automatically adjusting the rate of environmental perturbation to the intrinsic dynamics of the GRN under consideration without any prior knowledge about the architecture and dynamical feature of this GRN. This specific feature could be exploited in the future for ensuring the stability of microbial population in continuous cultivation, or to make the acquisition of fundamental knowledge about the role of biological noise in microbial response strategies to environmental fluctuations. The former perspective is in frame with a developing field of research i.e., cybergenetics. Cybergenetics uses computer interface for controlling in real-time biological processes [46][47]. This approach has been demonstrated to be very useful for controlling a variety of GRNs, including the antithetic motif [21][48] and the toggle switch [32].

## Acknowledgements

TMN is the recipient of a VIED PhD grant provided by the Vietnamese government. JAM and BZ are the recipient of a postdoctoral grant provided through two Era-Cobiotech projects, YBISCUS and ComRaDes respectively (funding through the European Union H2020 horizon program and Service Public de Wallonie (SPW)).

## Notes

https://gitlab.uliege.be/F.Delvigne/paper_segregostat_arabinose

### Competing Interest Statement

The authors have declared no competing interest.

